# Mapping Co-regulation Pathways among Ligand Binding sites in RyR1

**DOI:** 10.1101/676841

**Authors:** V. R Chirasani, K. I Popov, G. Meissner, N. V Dokholyan

## Abstract

Ryanodine receptor 1 (RyR1) is an intracellular calcium (Ca^2+^) release channel required for skeletal muscle contraction. RyR1 is co-regulated by multiple activators – Ca^2+^, ATP and caffeine (CFF), yet the mechanism of co-regulation and the action synergy of these activators is unknown. Here, we report the detailed network of allosteric connections between the three ligand sites and the pore region in (i) Ca^2+^ bound – closed, (ii) ATP/CFF bound – closed, (iii) Ca^2+^/ATP/CFF bound – closed and (iv) Ca^2+^/ATP/CFF bound – open RyR1 states. We find that two dominant paths mediate the communication between the Ca^2+^ binding site and pore region in Ca^2+^-only state. ATP/CFF-only occupied – closed RyR1 has two additional paths with CFF-but not ATP-occupied path using part of the closed Ca^2+^-only pathway. In the presence of Ca^2+^, ATP and CFF, major differences between the open and closed states are identified with both using part of the paths of the closed Ca^2+^-only and ATP/CFF-only states. We find that the three activators Ca^2+^, ATP, and CFF propagate their effects to the pore region through a network of partially overlapping pathways. Such coordination of allosteric pathway underlies the molecular basis of synergy of channel regulation by multiple activators.

**Statement of Significance:** RyRs are a group of Ca^2+^ channels that bind to several endogenous modulators and regulate Ca^2+^ release through closed-to-open gating transition. Despite the high-resolution structural data available for RyR1, the allosteric mechanism of RyR1 gating remains elusive. In this study, we employed graph-theoretical approach to demonstrate the allosteric network of synergistic interaction among various activators in RyR1. To our knowledge, for the first time we were able to identify the co-regulation among ligand sites in RyR1 to regulate the closed-to-open gating transition. The explored allosteric coupling in RyR1 may assist in designing advanced therapeutics for several debilitating diseases. Our findings in this study will assist to design new strategies for controlled allosteric regulation of RyR1 functionality in future.

## Introduction

RyRs are homo-tetrameric intracellular Ca^2+^ channels that account for the rapid release of Ca^2+^ ions from the sarcoplasmic reticulum (SR) into the cytoplasm, a critical event that prompts the contraction of muscle cells (1–5). RyRs are the largest known ion channels with molecular mass ∼2.2 megadaltons and are comprised of four identical monomers with each monomer containing approximately 5000 residues. There are three RyR isoforms in mammalian cells: RyR1 (skeletal muscle), RyR2 (heart muscle), and RyR3 (many tissues but in lower concentrations). RyR2 and RyR3 are activated by Ca^2+^-induced Ca^2+^ release (Calcium Induced Calcium Release, CICR), in which Ca^2+^ activates RyR through the release of Ca^2+^ ions. In contrast, RyR1 is controlled by Ca_v_1.1 voltage-gated Ca^2+^ channels, and Ca^2+^ by a mechanism not well understood (3, 6, 7).

Many channel endogenous modulators control RyR1 such as Ca^2+^, ATP, Mg^2+^ and calmodulin and exogenous effectors such as caffeine (CFF) and ryanodine (3). The binding sites of several RyR modulators are within 200 Å from the pore region in RyRs (8), which denotes the significance of allosteric communications within the channels. Some previous studies suggest the possibility of allosteric regulation; however, the mechanisms remain to be unraveled. More than eight-hundred mutations in RyR-isoforms have been reported to associate with several debilitating diseases such as central core disease and malignant hyperthermia in skeletal muscle, and catecholaminergic polymorphic ventricular tachycardia in cardiac muscle (9–12). While numerous clinical studies have been performed to emphasize the aberrant function of RyRs in disease mechanisms, the molecular events responsible for regulation of RyRs remain mostly unknown.

Recent advances in single-particle cryo-electron microscopy (cryo-EM) have captured RyR1 in multiple functional states and provided insights on RyR1 regulation by key activators Ca^2+^, ATP and CFF (8, 13–18). Particularly, the high resolution cryo-EM structures solved by Des Georges et al. (17) emphasized that the binding of Ca^2+^, ATP, and CFF can trigger local and global structural changes in RyR1 to induce channel opening. Mutagenesis and single-channel recordings revealed that two glutamate residues and one threonine residue in the Ca^2+^ binding site have a critical role in the Ca^2+^-dependent activation of RyR1 (19). Murayama et al. (20) showed that a tryptophan residue in the CFF-binding pocket of RyR1 and RyR2 regulates Ca^2+^ sensitivity by controlling the structure of the Ca^2+^ -binding site. Additionally, recent computational studies proposed that allosteric gating of RyR1 can be highly affected by extensive structural changes in peripheral and central domains of RyR1 (21, 22). While the cryo-EM structures and experimental studies provide valuable insights on ligand binding and functional states of RyR1, the allosteric mechanisms that couple and synergize communication between the ligand binding sites and pore region in RyR1 are not well understood.

In this study, we (i) propose the structural basis of allosteric communication between the Ca^2+^-binding site and pore region in open and closed RyR1 channels, (ii) show how binding of ATP and CFF can alter the allosteric network and affect Ca^2+^-free and Ca^2+^-occupied channel activities, and (iii) determine how conformational changes, related to channel opening or closing result in alteration of the allosteric network. We propose an allosteric network of synergistic interaction between the three activators Ca^2+^, ATP and CFF and how their interactions with RyR1 propagate their effects to the pore region through a network of connecting amino acid residues.

## Materials and Methods

### Mapping allosteric communications in different functional states of RyR1

We considered the cryo-EM structures of Ca^2+^ bound - closed RyR1 (PDB ID: 5T15), ATP/CFF bound - closed RyR1 (PDB ID: 5TAP), Ca^2+^/ATP/CFF bound - closed RyR1 (PDB ID: 5TAQ), and Ca^2+^/ATP/CFF bound - open RyR1 (PDB ID: 5TAL) and employed our recently developed methodology (23, 24) to delineate allosteric communication pathways between the ligands Ca^2+^, ATP, CFF and Gly-4934 on cytoplasmic extension of S6 (S6c) helix. Briefly, based on RyR1 tetrameric structure, we constructed a weighted graph S(N,E) – a mathematical object used to quantify relations between pairs of protein residues or ligands. The weighted graph S(N,E) is built by representing each residue as a node, n_i_, connected with other nodes of the graph by edges, e_ij_, where i,j=(1…p) and i≠j. The edges are weighted according to the number of contacts between heavy atoms in corresponding pair of amino acids of the protein or ligands. To choose crucial allosteric residues (nodes) we analyzed propagation of the communications through the network of nodes corresponding to the graph S(N,E). The product of edge weights along the paths was used as metrics to quantify communication strength along the pathway. To delineate allosteric paths between source and target nodes (either ligand–ligand or ligand–Gly-4934), we computed a series of rank-ordered optimal paths using Yen’s and Dijkstra’s algorithms (25, 26). Top 10,000 optimal paths were then clustered based on their similarity using hierarchical clustering algorithm. The following function were used as the paths dissimilarity measure:

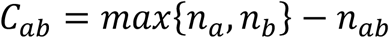

Where, *n*_*a*_, *n*_*b*_, and *n*_*ab*_ (0 ≤ *n*_*ab*_ ≤ *min*{*n*_*a*_,*n*_*b*_}) denotes number of nodes in path a, path b, and number of common nodes in paths a and b. It can be seen that *C*_*ab*_ would help to cluster pathways with small deviations (large *n*_*ab*_). After each step of clustering the pathways, we consider that the representative pathways carry the communication equal to the sum of those carried by merged pathways. There are several ways to choose the cutoff for hierarchical clustering (27). The detailed methodology on constructing a protein graph based on edge weights can be found in ref: (24). In this work, we found that the allosteric networks always had several strongest distinctive pathways that were carried through the clustering procedure and was always becoming representative pathways in the end of clustering. Based on this observation, we choose the cutoff based on the final number of representative pathways and analyzed only top five pathways (Figure 2 and Figures S1 – S6). We performed allosteric pathway analyses on tetrameric rather than monomeric conformations of RyR1 in different functional states as stated above. All pathway figures shown in this study were rendered using PyMOL (The PyMOL Molecular Graphics System, Version 2.0 Schrödinger, LLC).

**Figure 1.**
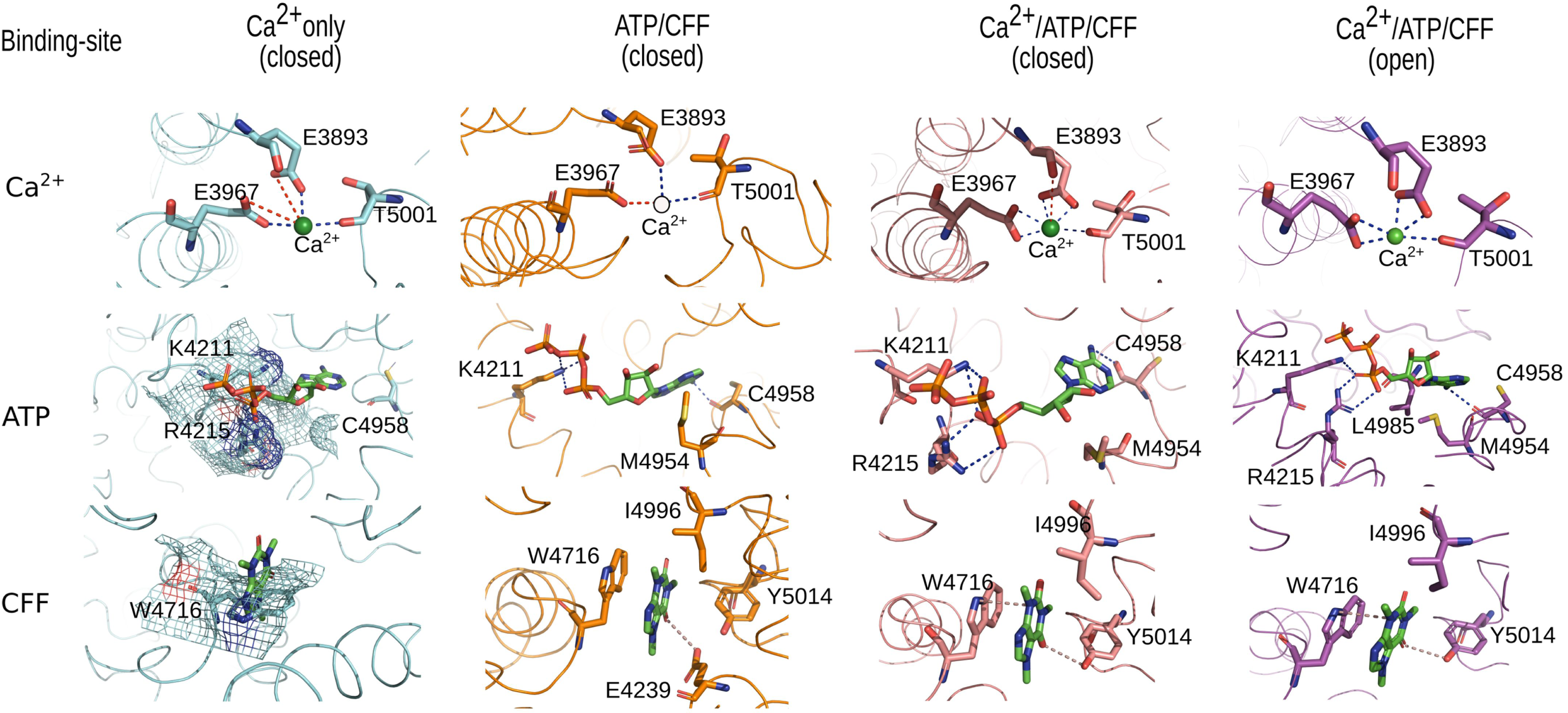
Interactions of Ca^2+^, ATP, and CFF in different functional states of RyR1s. Residues displaying electrostatic, van der Waals, and stacking interactions with Ca^2+^, ATP, and CFF in Ca^2+^ bound - closed RyR1 (PDB ID:5T15), ATP/CFF bound - closed RyR1 (PDB ID:5TAP), Ca^2+^/ATP/CFF bound - closed RyR1 (PDB ID:5TAQ) and Ca^2+^/ATP/CFF bound – open RyR1 (PDB ID:5TAL) are depicted in licorice representation. Backbone of RyR1 is shown as ribbon. Ca^2+^ is shown as a green sphere. ATP and CFF are shown in licorice representation. Residues exhibiting steric clashes with ligand are shown in mesh representation. Strong electrostatic interactions (<3.2 Å) are shown as blue dashed lines and weak electrostatic interactions (<4 Å) as red dashed lines. repulsive interactions (<3.5 Å) are depicted as dashed lines in salmon color.

**Figure 2.**
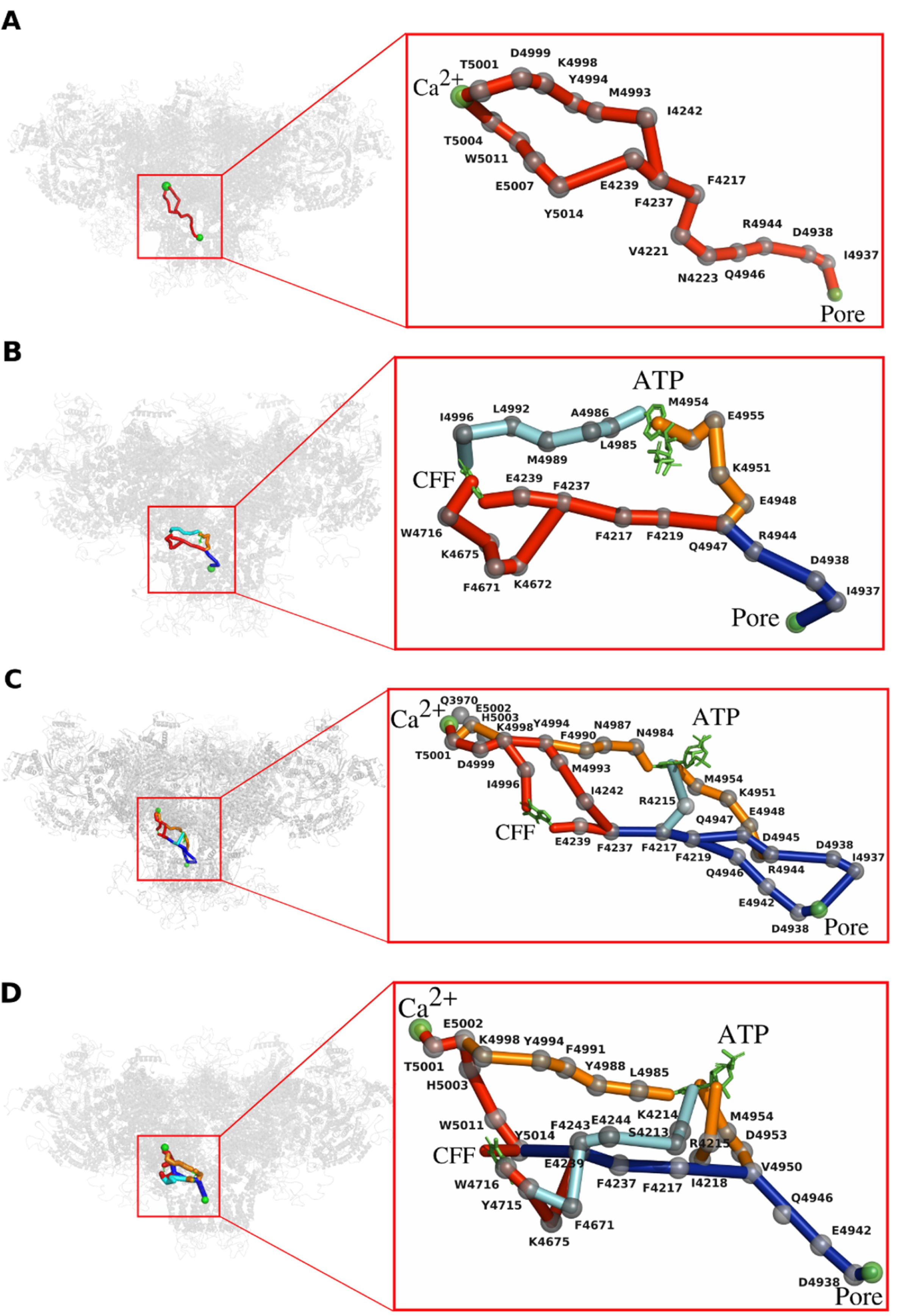
The detailed network of allosteric connections among the three ligand sites and the pore region in (a) Ca^2+^ bound - closed RyR1 structure (PDB ID: 5T15), (b) ATP/CFF bound - closed RyR1 structure (PDB ID: 5TAP), (c) Ca^2+^/ATP/CFF bound - closed RyR1 structure (PDB ID: 5TAQ), and (d) Ca^2+^/ATP/CFF bound - open RyR1 structure (PDB ID: 5TAL). Protein is shown in transparent gray cartoon representation, ATP and CFF are shown in green licorice, and Ca^2+^ and pore lining residue G4934 are shown in green spheres. Color code for allosteric pathways shown in this figure is as follows: Ca^2+^ – pore and Ca^2+^ – CFF – pore in red, Ca^2+^ – ATP – pore in orange, and CFF – ATP in cyan. Common allosteric pathways between any two regions are colored in blue. Cα atoms of allosteric information passing residues (hot spot residues) are shown in gray spheres.

### Single-channel measurements

Our single-channel measurements relied on inherent properties of RyR1, where RyR1 exhibits high conductance of K^+^ ions relative to Ca^2+^ and is impermeable to Cl^−^ ions. Channel activities were measured at pH 7.4 using 20 mM KHEPES, and 0.25 M KCl on both sides of the bilayer, 2 μM trans (SR luminal), and the denoted cis (cytosolic) additions. The filtered electrical signals at 2 kHz were digitized at 10 kHz and probed at 50% threshold setting (29). Data acquisition and analysis of 2-min recordings used pClamp (Axon Instruments).

## Results and Discussion

Recently published cryo-EM structures of RyR1 in multiple functional states have identified the binding sites for the channel activators Ca^2+^, CFF, and ATP at inter-domain interfaces of the C-terminal domain (8, 13–18, 30). Precisely, Ca^2+^ binding site is identified in the large cytoplasmic domain of RyR1, ATP-binding site is located close to pore region at the interface of two chains, and CFF binding pocket is located in relatively close proximity (∼20 Å) to Ca^2+^ binding site. Comparison of cryo-EM density maps obtained from reconstructions that have either only Ca^2+^ or only ATP/CFF deciphered considerable variations in densities of core of RyR1, designating synergistic activation of RyR1 by Ca^2+^ and ATP/CFF (17). Structurally, the co-localization of ATP-, CFF-, and Ca^2+^-binding sites at interfaces of the CTD might suggest possible allosteric communication between these binding sites in co-regulating channel opening and/or closing. However, the molecular details pertaining to the allosteric communication among these binding sites is far from complete.

To elucidate the structural basis and construct a possible allosteric network among Ca^2+^-, ATP-, and CFF-binding sites for channel activation, we performed allosteric pathway analysis on tetrameric conformations of Ca^2+^ bound - closed RyR1 structure (PDB ID: 5T15), ATP/CFF bound - closed RyR1 structure (PDB ID: 5TAP), Ca^2+^/ATP/CFF bound - closed RyR1 structure (PDB ID: 5TAQ), and Ca^2+^/ATP/CFF bound - open RyR1 structure (PDB ID: 5TAL). To delineate the allosteric pathways among different ligand binding sites and to check how the conformational changes in channels affects these communications, we employed graph-theoretical approach to depict allosteric communications between any chosen ligand sites in RyR1 structures. We analyzed relative information transmitted through each residue in a specific pathway and selected allosteric hot spot residues by evaluating residues conducting and transmitting maximum allosteric information. We generated bar charts specific to each pathway based on the amount of relative allosteric information transmitted by each residue specific to that pathway and subsequently highlighted allosteric hot spot residues crucial for communication (Figures S1 – S6). Thus, we not only identified crucial pathways between the ligand binding sites and the pore region, but also determined residues with maximal contribution to the allosteric couplings.

### Ca^2+^ to pore pathway propagates through CFF-binding site but evades ATP-binding site in Ca^2+^ bound - closed RyR1

The binding site of Ca^2+^ is constituted by the side chains of Glu-3893 and Glu-3967 (core solenoid) and the backbone of CTD residue Thr-5001 (Figure 1). It is apparent from cryo-EM data that the putative binding sites for ATP, CFF, and Ca^2+^ are situated in close proximity to each other and are strongly coupled to regulate the binding or release of Ca^2+^, either directly or indirectly (8, 13–18, 30). To delineate the influence of CFF- and ATP-binding sites on Ca^2+^ to pore communication, we first considered Ca^2+^ bound - closed RyR1 structure (PDB ID: 5T15) and employed our computational approach to understand the impact on allosteric signaling between Ca^2+^–pore regions in the absence of ATP- and CFF-binding. Analysis of our computationally identified allosteric network in Ca^2+^ bound - closed RyR1 structure allowed us to highlight two main Ca^2+^–pore pathways with several important communication residues (Figure 2(a) and Figure S1(a)). Interestingly, out of several allosteric pathways originated from Ca^2+^ binding site, the most prominent pathway initiated from Ca^2+^-binding pocket residue Thr-5001 and progressed through CTD residues Thr-5004, Tyr-5011, Glu-5007, and Tyr-5014. Then the allosteric communication persists in CFF-binding site mediated by CFF-pocket residues Tyr-5014 and Glu-4239. The electrostatic interactions between Glu-4239 and Phe-4237, aromatic interactions between Phe-4237 and Phe-4217, and hydrophobic interactions among Phe-4217, Val-4221, and Asn-4223 propagate the allosteric communication along CTD. Finally, the allosteric signal from Asn-4223 transport to the S6c helix residues through polar interactions among Gln-4946, Arg-4944, Asp-4938, and Ile-4937 residues in order to communicate with pore region. Hence, our analyses suggest that the communication between Ca^2+^ to pore is persistent through CTD and Thumb and Forefingers (TaF) domains, which has avoided the ATP-binding site explicitely.

The constricted ATP- and CFF-binding pockets in the absence of corresponding ligands prompted the evolution of a new communication pathway between Ca^2+^ to pore region. Specifically, a secondary communication pathway is initiated from Ca^2+^-binding pocket residue Thr-5001 that progressed along the CTD domain residues Asp-4999, Lys-4998, Tyr-4994, and Met-4993 through either covalent or hydrophobic interactions (Figure 2(a)). The allosteric signaling further advanced from CTD domain to TaF domain through tightly packed Met-4993 and Ile-4242 sidechains. The aromatic interactions between Phe-4237–Phe-4217 have subsequently propagated the allosteric communication along TaF domain. Finally, as observed in the primary pathway, allosteric signal from Phe-4217 transferred to S6c helix residues through polar interactions among Gln-4946, Arg-4944, Asp-4938, and Ile-4937 residues in order to communicate with pore region. *It is evident from our analysis that Ca*^*2+*^ *to pore communication in Ca*^*2+*^*-only occupied RyR1 propagates through CFF-binding site but evades ATP-binding site.* Thus, we can hypothesize that the binding of ATP may not critically influence Ca^2+^ sensitivity and sensitizing activity of Ca^2+^ on CFF in Ca^2+^ - only bound closed RyR1 system..

### CFF and ATP communicate the pore region through divergent and independent pathways in ATP/CFF bound - closed RyR1

Ca^2+^ ions play a crucial role in the regulation of RyRs through binding in micro-molar concentration at high affinity binding sites and in milli-molar concentrations at low affinity binding sites (1, 3). Additionally, the regulation of RyR1 is mediated by ATP and CFF, which strengthen Ca^2+^-gated RyR1 activities. However, understanding the effects of CFF and ATP binding on RyR1 gating in the absence of Ca^2+^ is far from complete. Here, we employed efficient computational techniques complemented by percolation theory to decipher allosteric signaling from CFF- and ATP-binding sites to pore region (PDB ID: 5TAP). Our allosteric pathway analysis on ATP/CFF bound - closed RyR1 denote that two sets of communications initiated from CFF-binding pocket residues Trp-4716 and Glu-4239 have propagated the allosteric signal from CFF binding site to the TaF domain (Figure 2(b)). The allosteric communication with in TaF domain from Glu-4239 to Phe-4237 is further advanced by non-covalent interactions. Conversely, the allosteric communication from Trp-4716 pass through Lys-4675, Phe-4671 and Lys-4672 to reach the TaF domain residue Phe-4237. The converged allosteric signal, mediated by aromatic interactions between Phe-4237–Phe-4217 and non-covalent or aromatic interactions Phe-4217–Phe-4219 and Phe-4219–Gln-4947 further progressed through TaF domain. Finally, the allosteric communication propagated from Phe-4219 to S6c helix residues through polar interactions among Arg-4944, Asp-4938, and Ile-4937 residues in order to couple with the pore region.

Similarly, the communication from ATP-binding site is initiated by the ATP-binding pocket residue Met-4954 and progressed through CTD residues Glu-4955, Lys-4951 and Glu-4948. Salt-bridge interaction between Glu-4948 and Arg-4944 has passed the allosteric signal initiated from ATP-binding site to the S6c helix, which further advanced through the S6c helix residues Gln-4947, Arg-4944, Asp-4938, and Ile-4937 before terminating at the pore region. Thus, our analysis identifies two independent communication sub-networks from CFF and ATP paths (Figure 2(b)) that converge in to one when they reach S6c helix. Our allosteric analysis displayed in few instances that the allosteric signal from CFF- or ATP-binding site passed from S6c helix in one monomer to S6c helix in another monomer before terminating at the pore region. We have also analyzed allosteric pathways between ATP- and CFF-binding sites to understand the co-regulation between these sites (Figure S4(a)). The allosteric communication between ATP- and CFF-binding sites progressed through CTD residues: Leu-4985, Ala-4986, Met-4989, Leu-4992, and Ile-4996, which are completely diverse compared to the residues involved in allosteric signaling from CFF- and ATP-binding sites to the pore region (Figure S2(a), S3(a)). Hence, the communication between CFF- and ATP-binding sites in the absence of Ca^2+^ binding is independent from ATP-pore and CFF-pore pathways. Thus, the discrete nature of pathways in CFF/ATP bound – closed RyR1 highlights the significance of Ca^2+^ binding for allosteric co-regulation among ATP-, CFF-, and pore regions.

### Numerous pathways exist in Ca^2+^/ATP/CFF bound – closed RyR1 structure to communicate between Ca^2+^ and pore regions

It was apparent from the alignment of closed and open RyR1 structures solved under Ca^2+^/ATP/CFF conditions that, RyR1 exhibits conformational variability in transmembrane region, which constricts the pore region and blocks the passage of a hydrated monovalent or divalent cations (17). Even in the presence of activator molecules, RyR1 (class III+IV) (Ca^2+^/ATP/CFF bound - closed RyR1 structure) exhibits closed conformation. Hence, we checked the inter communication between ligand sites in Ca^2+^/ATP/CFF bound - closed RyR1 structure through allosteric pathway analysis (Figure S1(b)). The allosteric connection between ATP- and CFF-binding sites is confined to TaF domain and is progressed through TaF domain residues Glu-4239 – Phe-4237 – Phe-4217 – Arg-4215 (Figure S4(b)). Furthermore, our allosteric pathway analysis identified that CFF-binding site significantly mediates the communication between Ca^2+^ binding site and pore region (Figure 2(c)). Specifically, the allosteric communication between Ca^2+^ to pore region is propagated through any of the following three pathways:

(i) Thr-5001 – Asp-4999 – Ile-4996 – **CFF** – Glu-4239 – Phe-4237 – Phe-4217 – Phe-4219 – Gln-4946 – Glu-4942 – Asp-4938 – Gly-4934

(ii) Thr-5001 – Asp-4999 – Lys-4998 – Tyr-4994 – Met-4993 – Ile-4242 – Phe-4237 – Phe-4217 – Gln-4947 – Arg-4944 – Asp-4938* – Ile-4937* – Gly-4934

(iii) Thr-5001 – Glu-5002 – His-5003 – Lys-4998 – Tyr-4994 – Phe-4990 – Asn-4987 – Asn-4984 – **ATP** – Met-4954 – Lys-4951 – Glu-4948 – Asp-4945 – Arg-4944 – Asp-4938* –Ile-4937* – Gly-4934

The allosteric signal in pathways-II and –III passed from S6c helix in one monomer to S6c helix in another monomer before terminating at the pore region (denoted by * above) (Figure 2(c)).

The residue interactions involved in above-mentioned pathways dominated mostly by polar and aromatic interactions. Furthermore, the partial alignment between CFF – ATP and Ca^2+^ – pore pathways in the vicinity of CFF binding site demonstrate the regulatory role played by CFF in optimizing Ca^2+^ to pore communication.

### CFF-binding site mediates the communication between Ca^2+^ to pore in Ca^2+^/ATP/CFF bound - open RyR1 structure

Recent cryo-EM studies on RyR1 by des Georges et al., furnished detailed structural information on Ca^2+^-, ATP-, and CFF-binding sites composition and location (Ca^2+^/ATP/CFF bound – open RyR1) (17). The CFF-binding site is located at the interface of S2S3 domain and CTD, approximately 25 Å from ATP-binding site, and is lined by residues Glu-4239, Phe-3753, Trp-4716, Ile-4996, and Tyr-5014 (Figure 1). The ATP-binding site is located at the S6c – CTD interface with Lys-4211, Lys-4214, Arg-4215, Met-4954, Glu-4955, Phe-4959, Thr-4979, and Leu-4985 as binding site residues (Figure 1). Binding of CFF alter the orientation of Trp-4716 and induce conformational changes in CTD to shrink Ca^2+^ binding pocket for more favorable binding of Ca^2+^ (20). The binding of Ca^2+^, ATP, and CFF at respective binding sites initiates reorientation of the CTD and induces flexible alterations in the activation module. The activation module, thus acquires a conformation and orientation responsible for pore enlargement. Hence, the extent of channel opening probability is largely dependent on the binding of multiple activators (Ca^2+^, ATP, and CFF) at different sites on the CTD. Recent study on RyR1 using single-channel measurements complemented by computational analysis emphasized strong coupling between S6 pore-lining helix and Ca^2+^ permeation (31). As these major structural rearrangements in the open RyR1 channel can completely reorganize allosteric communication network compare to the closed state, we performed allosteric pathway analysis on RyR1 structure in its new functional state.

Cryo-EM data suggests that Ca^2+^ has strong electrostatic interactions with three conserved amino acid residues (Glu-3893, Glu-3967, Thr-5001) in Ca^2+^ binding pocket (Figure 1) (17). Our allosteric analysis suggests that the CICR activity initiates from one of the Ca^2+^ binding pocket residues, Thr-5001 and propagates through the neighbor residues Glu-5002, and His-5003. The strong aromatic interactions between His-5003 – Trp-5011 and Trp-5011 – Tyr-5014 allow the progression of allosteric communication from His-5003 to CFF-binding pocket residue Tyr-5014. Here, CFF-binding site residues Tyr-5014 and Glu-4239 play significant role in propagating communication between Ca^2+^ and pore regions by transferring the allosteric signal to TaF residues Phe-4237 and Phe-4217. Thus, we can assume that the allosteric communication during CICR activity may be modulated by CFF molecule. Subsequent progression of allosteric signal from Ca^2+^ binding site to pore region passes through Phe-4237 and Phe-4217 residues supported by strong aromatic interactions. Finally, the communication received from Phe-4217 pass through the residues of S6c helix, i.e., Val-4950, Gln-4946, Glu-4942, and Asp-4938 before reaching the pore region. We have also observed two secondary allosteric communication pathways between Ca^2+^ and pore regions mediated by ATP-binding site residues Met-4954 and Arg-4215 (Figure 2(d) and Figure S1(c)). Additionally, some pathways pass through S6c helix of the other monomer before coming back to the S6c helix of the same monomer during communication with the pore region for CICR activity.

Since (a) Ca^2+^ binding site and pore region are present on either side of the vector passing through ATP- and CFF-binding sites, and (b) Ca^2+^ is distal from CFF–ATP–pore inter-communication network, the communication pathways from CFF- and ATP- to pore region avoid Ca^2+^ binding site (Figure S2(c) and Figure S3(c)). This observation suggests that the binding of Ca^2+^ does not have direct impact on ATP – CFF communication. Cryo-EM data indicate that CFF-binding site comprised of five conserved residues (Phe-3753, Glu-4239, Trp-4716, Ile-4996, and Tyr-5014) where CFF has aromatic interactions with Trp-4716, polar interactions with Glu-4239, Phe-3753, and Tyr-5014, and hydrophobic interactions with Ile-4996 (Figure 1). Our allosteric pathway analysis suggests that the allosteric communication from CFF-binding most likely propagates through either Tyr-5014 or Trp-4716 to the TaF domain by one of the following two routes:

(i) Trp-4716 – Tyr-4715 – Lys-4675 – Glu-4239 – Phe-4237 – Phe-4217;

(ii) Tyr-5014 – Glu-4239 – Phe-4237 – Phe-4217.

The type of interactions in both pathways are contributed mostly by polar, aromatic, and hydrophobic interactions. Once the allosteric communication from CFF-binding site reaches the TaF-domain, the allosteric signal subsequently progresses through Val-4950, Gln-4946, Glu-4942, and Asp-4938 to reach the pore region similar to Ca^2+^ – pore pathway. This observation suggests that communication between Ca^2+^ to pore region is mediated by CFF-binding site. Since Ca^2+^ to pore communication pathways mostly avoid ATP binding side, binding of ATP should not directly influence the primary communication between either Ca^2+^ to pore or CFF to pore regions. However, the fact that both these communications merge with the allosteric pathways going from ATP to the pore on the helix S6 might indicate that binding of ATP still modulates the ion channel gating control by affecting the final stage of the communications. Furthermore, the persistence of ATP – CFF signaling within TaF domain denotes strong correlation between ATP and CFF binding sites (Figure 2(d) and Figure S4(c)). Thus, we propose that the primary allosteric communication between Ca^2+^ to pore is predominantly regulated by CFF-binding site in Ca^2+^/ATP/CFF bound - open RyR1 structure, although there exist less probable pathways that propagate through ATP-binding site (Figure 2(d) and Figure S3). Hence, we conclude that the allosteric communication among Ca^2+^-, CFF-, and ATP-binding sites render significant role in RyR1 channel regulation. Although few additional residues participated in CFF – pore (Figure S2) and ATP – pore pathways (Figure S3), the allosteric communications in Ca^2+^/ATP/CFF bound - closed and open RyR1 structures are significantly similar.

## Conclusions

Allosteric transition, because of ligand binding, is a characteristic feature of proteins. Conformational changes in proteins are important to regulate protein function and other key biological processes. In this study, we analyzed the allosteric pathways between ligand binding sites and pore region of RyR1 in four functional states. We found that the opening or closing of pore region is allosterically coupled to Ca^2+^-, ATP-, and CFF-binding sites. Our results also depicted that the communication between Ca^2+^ to pore region is mediated by CFF-binding site in both open and closed RyR1 structures. Aromatic residues such as Trp-4716 and Tyr-4715 were identified to be a part of CFF-binding site that regulate the allosteric communication between Ca^2+^ to pore region, specifically in ATP/CFF bound – closed, Ca^2+^/ATP/CFF bound – closed, and Ca^2+^/ATP/CFF bound – open RyR1 structures (Figure 2(b), 2(c), and 2(d)). Previous experimental studies showed that the binding of CFF alter the orientation of Trp-4716, which induce conformational changes in CTD to modulate Ca^2+^ binding pocket for more favorable binding of Ca^2+^ (20). The subsequent binding of Ca^2+^ and ATP at respective binding sites initiates reorientation of the CTD and there by induces flexible alterations in the activation module. The activation module, thus acquires a conformation and orientation responsible for pore enlargement as evident from the cryo-EM data of Ca^2+^/ATP/CFF bound – open RyR1 structure. However, ATP-binding site does not seem to influence the primary communication between either of Ca^2+^ to pore and CFF to pore regions. Furthermore, the absence of Ca^2+^ has significantly altered CFF – ATP and CFF – pore pathways in ATP/CFF bound – closed RyR1 structure (Figure 2(b)) in comparison to respective pathways in other RyR1 functional states (Figure 2(c) and 2(d)). However, ATP – pore pathway was least affected by the presence or absence of Ca^2+^. Hence, allosteric communications either initiated from or transmitted through CFF-binding site were critically influenced by the binding or unbinding of other ligands. The comparison of allosteric pathways among different RyR1 functional states also denoted that the structural conformation of pore region (pore opening vs pore closure) significantly affected ATP pore and CFF – pore communications. Specifically, closed RyR1 structures (Figure 2(b) and 2(c)) showed irregular information exchange among ATP, CFF, and pore regions in comparison to the organized trade-off in open RyR1 (Figure 2(d)). In addition, CFF – Ca^2+^ pathway exhibited diverse changes and ATP – Ca^2+^ pathway displayed identical pattern in RyR1 – closed (Figure 2(c)) and RyR1 – open structures (Figure 2(d)). Thus, the allosteric network of synergistic interactions among various activators in RyR1 regulate the opening/closing of RyR1 pore region.

We identified unique residues specific to allosteric interactions in a specific functional RyR1 state. For example, TaF domain residues Val-4221 and Asn-4223 were identified as part of allosteric network in Ca^2+^ bound – closed RyR1 (Figure 2(a)). We have also identified some residues important for allosteric communications in Ca^2+^ bound – closed RyR1 and ATP/CFF bound – closed RyR1 structures, those are shared by allosteric pathways in Ca^2+^/ATP/CFF bound closed and open RyR1 structures. Specifically, Phe-4217, Phe-3237, and Phe-4239 from TaF domain; Ile-4937, Asp-4938, Glu-4942, Arg-4944, and Gln-4946 from pore S6c helix were spotted to participate actively in various allosteric interactions across different RyR1 functional states (Figure 2). We have further identified residues crucial for allosteric communication among Ca^2+^-, ATP-, CFF-binding sites, and pore region. Mutations in these residues severely affected RyR1 functionality. For example, T5001A have potentially altered channel opening properties (19). Interestingly, single channel measurements on RyR1 with pore-lining helix residue G4934 mutated to either Ala or Val depicted abnormal channel characteristics (31). Specifically, RyR1-G4934A variant displayed reduced ion selectivity, K^+^ conductance, and CFF-induced Ca^2+^release in comparison to RyR1-WT (31). Mutations in Ca^2+^ activation – pore pathway residues such as D4938N and D4945N, significantly diminished ion conductance and selectivity for Ca^2+^ in HEK293 cells (29). Furthermore, three single mutations: Q4947N, Q4947T, and Q4947S in S6c helix displayed reduced channel opening probability (P_o_) in comparison to RyR1-WT (32). Single RyR1-WT channel activities determined in the presence of the three channel activators Ca^2+^, ATP and caffeine deciphered that the addition of 2 mM ATP and 5 mM caffeine to cytosolic 30 µM Ca^2+^ significantly increased channel opening probability (P_o_) (Figure 3) (19). Thus, the correlation between loss of CFF sensitivity – response, Ca^2+^ activation upon mutating pore-lining helix residues, and enhanced channel opening probability (P_o_) in presence of ATP, CFF, and Ca^2+^ substantiates strong allosteric coupling among Ca^2+^, CFF, ATP, and pore regions (Figure 4). Due to constricted nature of the transmembrane region, several additional residues were involved in diverse allosteric communications between ligand sites in Ca^2+^/ATP/CFF bound - closed RyR1 structure (Figure 2(c)) in comparison to that of Ca^2+^/ATP/CFF bound – open RyR1 (Figure 2(d)). We have also identified in the allosteric network of Ca^2+^/ATP/CFF bound – closed RyR1 that there exists an intermediate pathway between Ca^2+^ and pore regions avoiding both CFF- and ATP- binding sites (Figure 2(c)).

**Figure 3.**
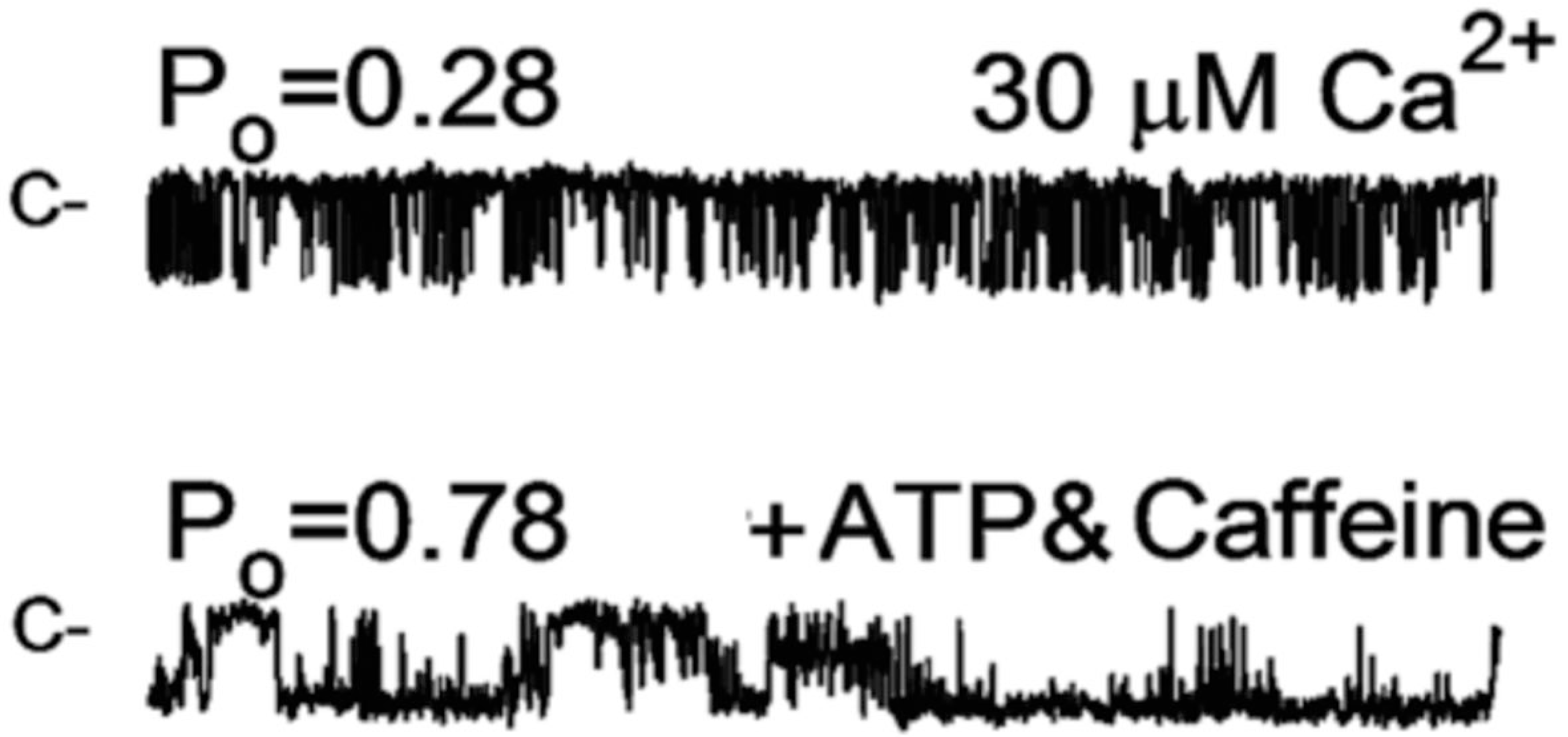
Effects of CFF and ATP on RyR1-WT channel open probabilities. Representative single channel currents are shown at −20 mV as downward deflections from the closed state (c--) in symmetrical 0.25 M KCl. Free cytosolic Ca^2+^ was 30 µM (upper trace) and 5 µM after the addition of 5 mM CFF and 2 mM ATP (lower trace). Modified from Figure 6A in ref. (19).

**Figure 4.**
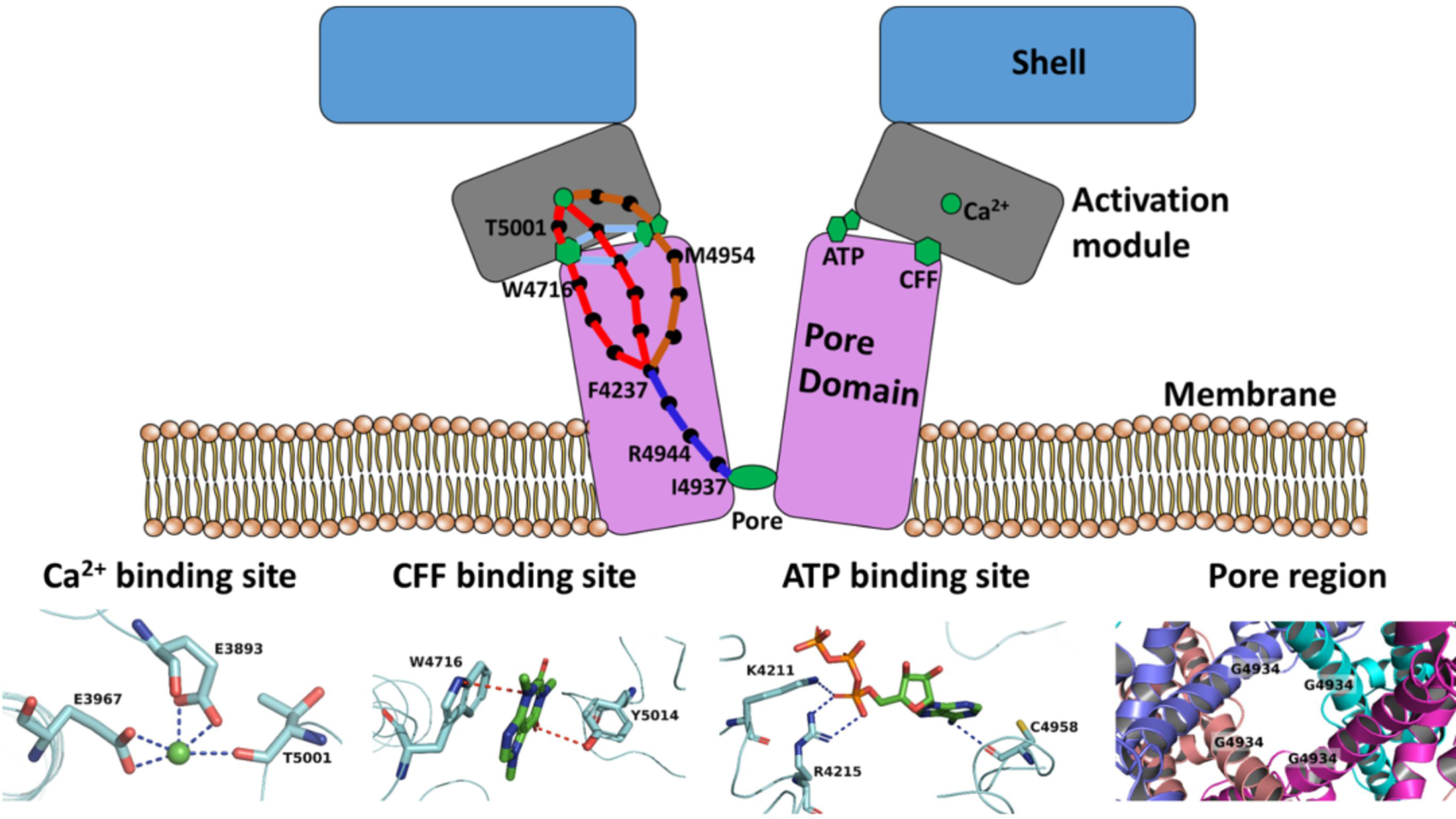
Molecular basis of synergy of channel regulation by multiple activators in RyR1. Schematic representation of residues responsible for initiating allosteric signal from each ligand-binding site and residues that modulate and propagate the allosteric communication to the pore region. Interactions of Ca^2+^, CFF, and ATP in respective binding sites and the organization of pore region in open RyR1 structure also shown for better understanding.

Finally, using our graph-theoretical approach, we demonstrated the allosteric network of synergistic interaction among various activators in RyR1 to control the opening/closing of RyR1 pore region (Figure 4). The allosteric coupling among ligand-sites and pore region explored in the current study may assist in designing advanced therapeutics for several debilitating diseases such as central core disease, catecholaminergic polymorphic ventricular tachycardia (CPVT), malignant hyperthermia, and heart related complications. Mutagenesis studies on unique and common allosteric residues among various RyR1 functional states explored in this study might help us to devise new methodologies for controlled regulation of RyR1 functions. Furthermore, this study suggests a possibility to design complex regulation of protein functionality by multiple allosteric activators.

## Supporting material

Supporting figures S1-S6 are freely available in the web.

## Author contributions

Conceived and designed the experiments: VRC, NVD, and GM. Performed the experiments: VRC and KIP. Analyzed the data: VRC. Contributed reagents/ materials/analysis tools: NVD and GM. Wrote the paper: VRC, KIP, NVD, and GM.

## Acknowledgements

We thank Dr. Jian Wang for developing an interactive web-server to depict graph-inspired pathways in RyR1. N.V.D. acknowledges the NIH support grants R01GM114015 and R01GM123247. N.V.D. and G.M. are supported by the NIH grant AR018687.

